# ERnet: a tool for the semantic segmentation and quantitative analysis of endoplasmic reticulum topology for video-rate super-resolution imaging

**DOI:** 10.1101/2022.05.17.492189

**Authors:** Meng Lu, Charles N. Christensen, Jana M. Weber, Tasuku Konno, Nino F. Läubli, Katharina M. Scherer, Edward Avezov, Pietro Lio, Alexei A. Lapkin, Gabriele S. Kaminski Schierle, Clemens F. Kaminski

## Abstract

The topology of endoplasmic reticulum (ER) network is highly regulated by various cellular and environmental stimuli and affects major functions such as protein quality control and the cell’s response to metabolic changes. The ability to quantify the dynamical changes of the ER structures in response to cellular perturbations is crucial for the development of novel therapeutic approaches against ER associated diseases, such as hereditary spastic paraplegias and Niemann Pick Disease type C. However, the rapid movement and small spatial dimension of ER networks make this task challenging. Here, we combine video-rate super-resolution imaging with a state-of-the-art semantic segmentation method capable of automatically classifying sheet and tubular ER domains inside individual cells. Data are skeletonised and represented by connectivity graphs to enable the precise and efficient quantification and comparison of the network connectivity from different complex ER phenotypes. The method, called ERnet, is powered by a Vision Transformer architecture, and integrates multi-head self-attention and channel attention into the model for adaptive weighting of frames in the time domain. We validated the performance of ERnet by measuring different ER morphology changes in response to genetic or metabolic manipulations. Finally, as a means to test the applicability and versatility of ERnet, we showed that ERnet can be applied to images from different cell types and also taken from different imaging setups. Our method can be deployed in an automatic, high-throughput, and unbiased fashion to identify subtle changes in cellular phenotypes that can be used as potential diagnostics for propensity to ER mediated disease, for disease progression, and for response to therapy.

## Introduction

The endoplasmic reticulum (ER) is the largest membranous structure in eukaryotic cells and acts as a platform for protein synthesis and quality control and for various organelle-interactions (Schwartz and Blower 2016). The healthy function of the ER depends on its dynamics and structure (Westrate et al., 2015), which are highly regulated by intra- and extracellular stimuli. The ER consists of distinct domains including sheets and tubules, and features growth tips and tubular connections, so called three-way junctions. Perturbations to the ER structure and dynamics caused by genetic defects or metabolic stress have been associated with a variety of diseases (Schönthal 2012), such as spastic paraplegias (HSPs) and Niemann Pick Disease type C (NPC). Hence, to understand the role of ER in diseases, it is important and necessary to characterise ER morphology comprehensively, which may provide powerful phenotypes to screen drugs against ER associated disorders. However, given the extent of the ER network and its complexity, the precise and quantitative measurement of ER topology and movement has remained challenging. The ER network in a single cell consists of thousands of interconnected tubules that undergo constant rearrangements *via* processes including continuous tubular elongation, contraction, and fusion. Furthermore, there are rapid transitions between sheet and tubular domains with distinct putative functions (Lu et al., 2020). Recently, capabilities have emerged to reveal such dynamic changes in ER topology in live cells, at sub-wavelength resolution. Structured illumination microscopy (SIM), for example, can be used to resolve details of ER topology and its rapid remodelling process (Nixon-Abell et al, 2016; Guo et al., 2018). However, the data have only been interpreted qualitatively, without attempts to quantify ER topology or its structural changes precisely. So far, no suitable metrics exist, nor analysis tools, that can be used for such a purpose. Compared to other organelles, such as mitochondria and lysosomes, which are structurally simpler organelles that are often well separated from one another, the ER consists of highly convoluted and structurally connected domains. The task is further complicated by the fact that the signal to noise ratio of images obtained during live cell microscopy is often poor, while a clear differentiation of the organelle from its background is required to ensure successful segmentation into tubular and sheet domains. For moving structures, and time lapse imaging, this becomes a formidable task.

A number of machine-learning based methods have been developed for the segmentation of cells (Stringer et al., 2021), mitochondria (Fischer et al., 2020; Lefebvre et al., 2021), and nuclei (Hollandi et al., 2020), which provide robust and precise classification of cell structures. However, to date, thresholding remains the standard method of use for ER segmentation (English and Voeltz 2013; Pain et al., 2019; Garcia-Pardo et al., 2021), a method which lacks both sensitivity and specificity and thus quantitative conclusions are hard to draw, especially in situations where image quality is compromised by noise. Alternative methods are based on labour intensive manual labelling of image data to generate specialised datasets for training of machine learning algorithms. These approaches do not generalise well to work with changing experimental setups or varying sample types (Extended Data Fig. 1) (Arganda-Carreras et al., 2017). An additional challenge for ER segmentation can be seen in temporal consistency. Conventional segmentation is performed on a frame-by-frame basis, and segmented structures in sequential (time-lapse) images lose temporal continuity and thereby cause artefacts (Belthangady and Royer 2019). Currently, there is no ER segmentation method capable of taking dynamic, spatial and temporal topology changes into consideration. Hence, more efficient and accurate classification schemes need to be developed for sequential imaging data, to be able to study ER structural changes as the they occur in live cells.

To address these difficulties, we developed ERnet, a deep learning-based software that automatically segments ER, classifies its domains into tubules and sheets, and quantifies structural and dynamic features in super-resolution image sequences obtained from live cells. We provided ERnet with an intuitive user interface to make it a broadly accessible tool for biologists (Extended Data Fig. 2) and to promote ER-related research in basic science and clinical applications. While conventional segmentation methods based on thresholding classify objects according to image intensity, ERnet is trained with large image datasets to model the domain knowledge of ER structures, *i.e*., the shapes of tubules and sheets. As a result, it enables feature specific segmentation with enhanced robustness, specificity, and sensitivity regardless of the pixel intensity in the images. After segmentation, ERnet quantifies topological features of the ER and recognises subtle changes in the ER structure and dynamics for various stress conditions, including gene knockout /knockdown, ATP depletion and Calcium depletion etc. To validate the method, we tested the segmentation accuracy of ERnet on *in vitro* models subjected to different genetic and metabolic manipulations, including cells mimicking phenotypes of HSP and NPC. Two phenotypes were identified as sensitive readouts of the ER response in these models, namely the degree of fragmentation of ER networks and the heterogeneity in tubule connections. Both are indicators for the functional state of the ER network, and can be used, *e.g*., to quantify the degree of disorganisation, shrinkage, and collapse of ER structures in models of disease. In summary, ERnet enables automated segmentation of ER structures and parametric analysis of ER topology in models used for genetic or therapeutic screening.

## Results

### The ERnet model architecture is optimised to segment and capture network information obtained from video-rate super-resolution imaging data

The general design of ERnet is schematised in Fig. 1a. First, the reconstructed sequential images of the ER were segmented in ERnet, followed by the classification of ER structures into tubules and sheets. The tubular structure was further skeletonised using a surface axis thinning algorithm (Lee et al., 1994). After this, the nodes and edges of the skeletonised ER were identified to plot a topology graph *via* a graph theory-based module (Peixoto, 2014). Instead of relying on the commonly applied convolution neural networks (CNN), our model builds upon a Vision Transformer architecture (Dosovitskiy et al., 2020) which outperforms a comparable state-of-the-art CNN with higher classification accuracy and four times fewer computational resources. Key to our method is that, rather than paying attention to the physical locations of the nodes, it focuses on the ER’s network features, *e.g*. the connectivity between nodes. For instance, metrics such as number of fragments and clustering coefficients can be extracted to determine the ER topology.

**Fig. 1:**
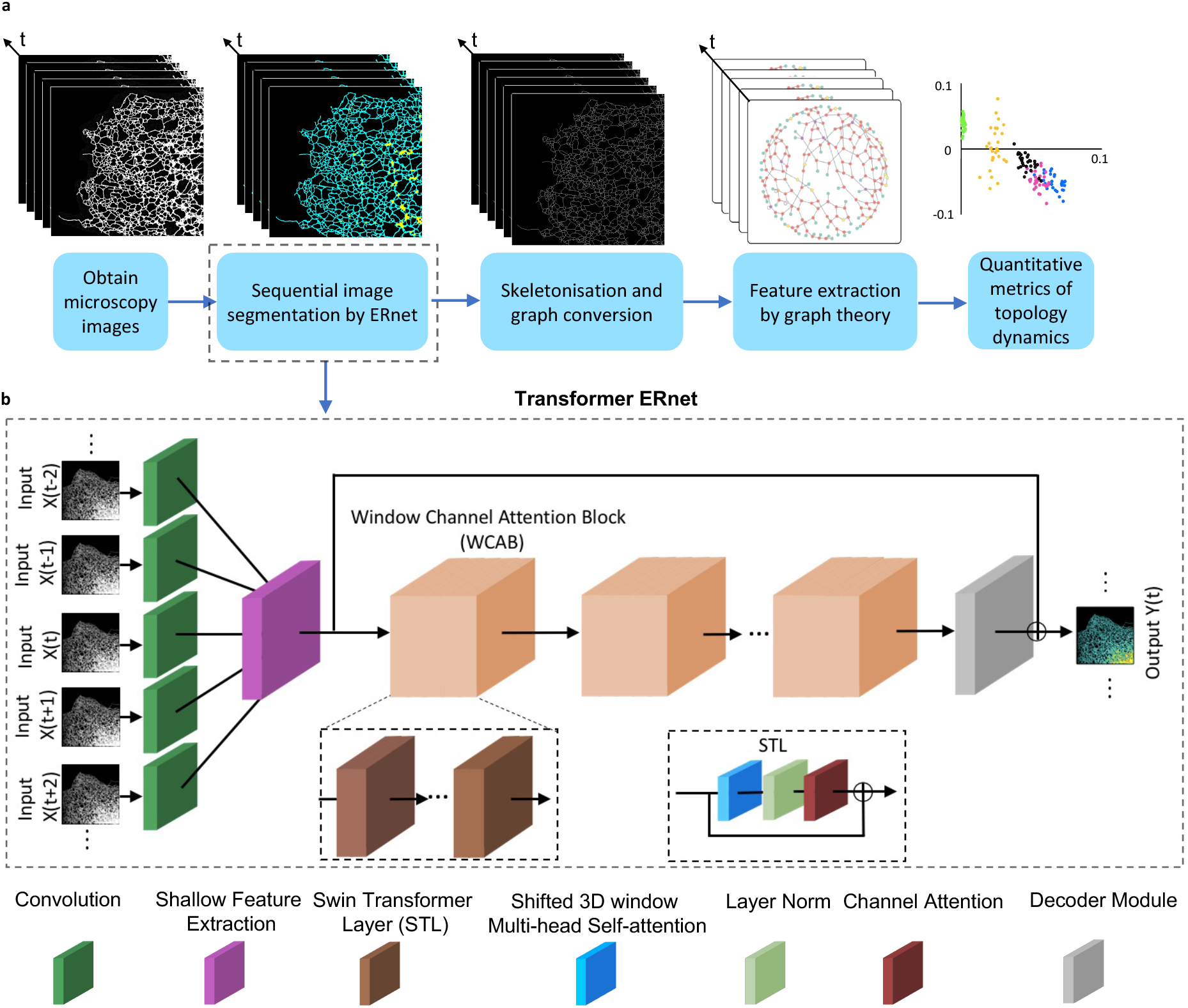
Workflow of ER structure segmentation and ERnet construction. a. The processing pipeline of ER segmentation and analysis. Time-lapse SIM images were first segmented by ERnet to classify the tubules and sheets. The tubular network of ER after segmentation was further skeletonised and the nodes and edges were identified to plot the connectivity graph. Using graph theory-based methods, we quantified the metrics of the ER network features that describe the topology and dynamics. b. The Transformer based architecture of ERnet. A moving window loads adjacent frames (X_t-*2*_ to X_t+*2*_) as inputs from the time-lapse images into ERnet. A shallow feature extraction module then projects the input into a feature map which is followed by a sequence of residual blocks denoted with Window Channel Attention Block (WCAB). Inside each WCAB, there is a sequence of Swin Transformer Layers (STLs).

The core component in our workflow is a Vision Transformer based model ERnet that performs the segmentation of the super-resolution images recorded at video rates (Fig. 1b). ERnet is designed to have a temporal window of five adjacent frames as input which permits the model to process sequentially correlated ER structures. By introducing a set of sequential frames with temporally overlapping structures, moving objects demonstrate a higher correlation than random background noise which improves the recognition of ER structures and allows the model to obtain more comprehensive domain knowledge that is critical to assess the structural integrity of the ER network correctly. To reduce the computational cost associated with the large data volumes generated by time sequenced imaging data, ERnet makes use of a so called 3D shifted window (Liu et al. 2021) that not only applies self-attention to information within specific individual images themselves but also to features that persist between different frames in the sequence. We also combine the multi-head self-attention (MSA) mechanism (Vaswani et al, 2017) with a channel attention mechanism (Christensen et al., 2022) in the ERnet, a design which makes the method more adaptive to different ER phenotypes.

### ERnet performs precise segmentation and topological analysis of the ER structures in sequential SIM images

The ER is a highly dynamic structure and at any instance thousands of tubules move and change position, direction, and network connections. The purpose of ERnet is to obtain quantitative information from the above ER structural changes which are closely linked to disease phenotypes. To quantify these intracellular changes, we first tested performance of ERnet using SIM images of COS-7 cells. Fig. 2a shows a single frame of the ER (grey) from a set of sequential images captured from a COS-7 cell expressing mEmerald-Sec61b (Nixon-Abell et al., 2016). The performed segmentation successfully identified the whole ER structure, differentiated it from the cytosol background and further classified it into tubular (cyan) and sheet domains (yellow) (Fig. 2a). Then, the tubular ER was skeletonised from the whole structure and the nodes (tubule junctions, shown in red) and edges (tubules, green) were identified as two key topological components to map the network connectivity *via* the Python package Graph-tool (Peixoto 2014).

**Fig. 2:**
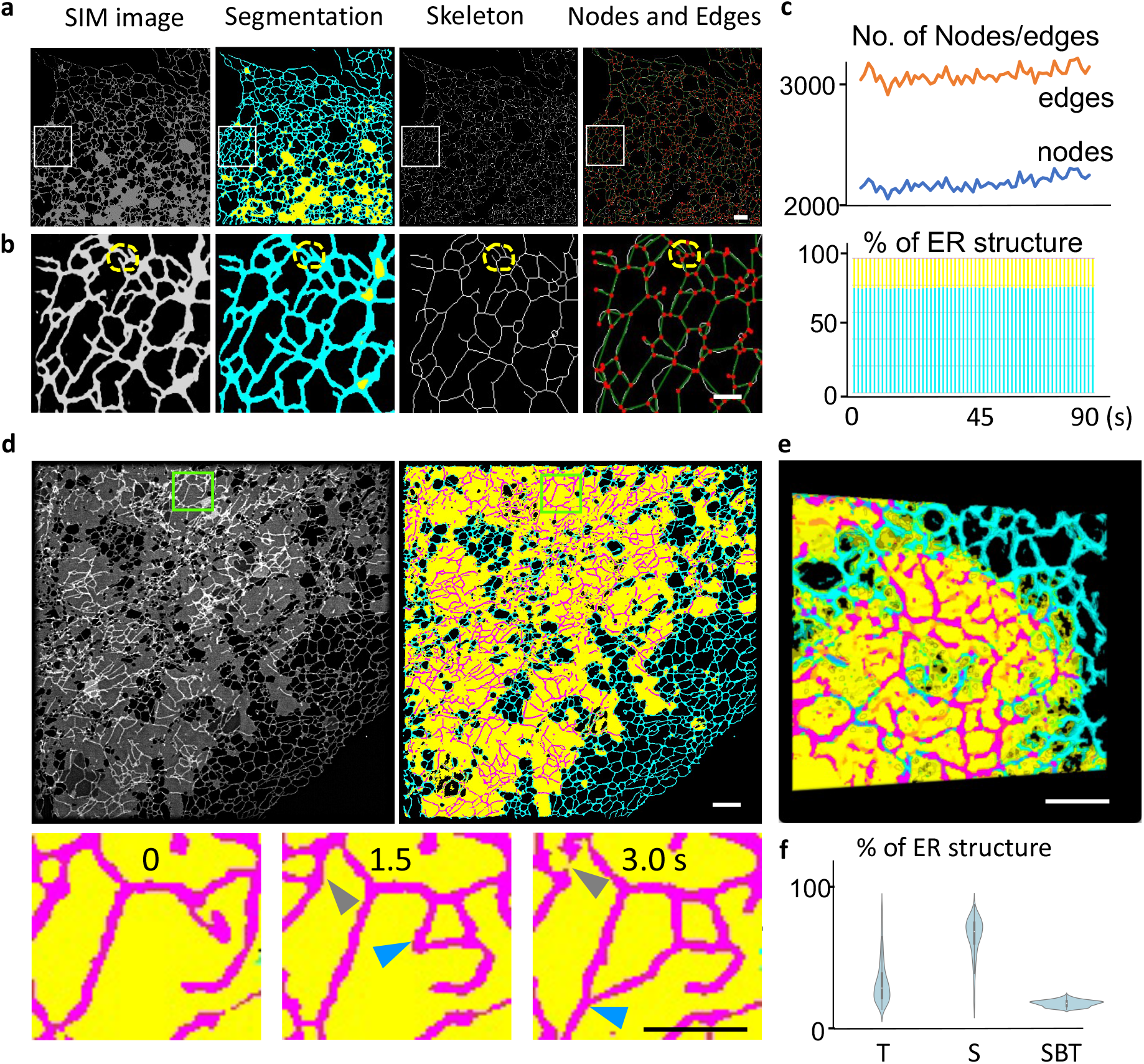
Semantic segmentation of ER and classification of tubules and sheets. a. An example of a segmentation result from video-rate SIM images of the ER. From left to right: 1) SIM image, 2) segmentation of ER tubular (cyan), sheet (yellow) and sheet-based tubule (magenta) region, 3) skeletonisation of the tubular domain, and 4) identification of nodes (red spots) and edges (green lines) based on the skeleton structure. Scale bar: 5 μm. b. Zoomed-in regions of the above panel. The yellow dashed circles indicate nodes that are closely positioned but can still be identified by ERnet. Scale bar: 2 μm. c. Quantitative analysis of the ER shown in (a). Top panel: quantification of edges and nodes of the ER tubules of the time-lapse frames. Bottom panel: percentage of the ER tubules (cyan) and sheet (yellow) of the time-lapse frames. See Source Data Fig. 2c. d. A representative frame from time-lapse images shows the structure of sheet-based tubules. Top left panel: a SIM image of the ER structure. Top right panel: segmentation of the three ER structures: sheet-based tubules (magenta), sheet (yellow), tubules (cyan). Bottom panel: three sequential frames showing the dynamic reshaping of sheet-based tubules from the above green boxed region. Blue arrows indicate a continuously elongating sheet-based tubule and grey arrows indicate a retraction. Scale bars: 5 μm (top) and 2 μm (bottom). See Source Data Fig. 2d for quantitative analysis. e. Volumetric view of 3D reconstruction of the sectioning SIM showing that the sheet-based tubules (magenta) are embedded in sheet domains (yellow). Scale bar: 2 μm (bottom). f. Violin plots of the percentages of tubules (T), sheets (S) and sheet-based tubules (SBT) in COS-7 cells (*N*=500), showing that the presence of the sheet-based tubules is a common feature of the ER network. See Source Data Fig. 2f.

SIM provide high spatial-temporal resolution of ER structures thus suitable for live cell imaging (Extended Data Fig. 3). A single pixel on the camera frame has a length scale of 42 nm in real space, almost a quarter of the average width of an ER tubule (~160 nm, measured as the average width on SIM images taken). This means that misclassification of a few, or even just one, image pixels can mean the difference between identification of a tubule as connected, or as disrupted. This leads to errors in the classification of network features, and *vice versa* to a bias when quantifying the network connectivity. In disease models, this could lead to erroneous phenotypes. The semantic segmentation of individual pixels from SIM images ensures the structural integrity of networks identified and prevents information loss, an improvement of traditional algorithms used in the past. Figs. 2a and b show how the method performs. A clear segmentation of ER structure (Fig. 2b) is achieved in regions containing dense ER tubule networks, as can be seen from the enlarged region indicated by the white box in Fig. 2a. This permits the distinction of tubules and their junctions in confined regions, measuring less than 300 nm across (highlighted by yellow dashed lines) with good structural detail. The segmented ER was then skeletonised (middle panel of Fig. 2a and b) and classified into edges (green tubules, right panel, Figs. 2a and b) and nodes (red spots, right panel, Figs. 2a and b). Finally, ERnet quantified the number of edges and nodes (top plot, Fig. 2c) and the percentage of areas covered by tubules and sheets (bottom plot, Fig. 2c), respectively, across the whole ER. Here, ER tubules were defined as linear branched structures and sheets as flat membrane cisternae as shown in Fig. 2a and d. Morphological features, such as the percentage of tubules/sheets among the whole ER, reflect ER status (Lu et al., 2020) and provide indications for possible ER defects. ER stress induced by an absence of the GTPase Rab7, which is known to modulate lysosome-ER contact sites, leads to the enlargement of ER sheets and the reduction of tubular domains in the cell periphery (Mateus et al., 2018). On the other hand, a depletion of protrudin, an ER reshaping protein, induces HSP associated ER dysfunctions by disrupting the sheet-to-tubule balance (Chang et al, 2013). Therefore, and as investigated in more detail in the subsequent sections, it is expected that the topological features of the ER, such as its connectivity, assortativity, or clustering coefficients, change for different phenotypes and with disease progression. It is worth highlighting that, although the ER tubular network underwent stark morphology changes (Movie 1) and demonstrated fluctuations in the numbers of nodes and edges (top panel, Fig. 2c) within individual recordings, its tubule and sheet percentage among the whole ER remained stable (bottom panel, Fig. 2c), which suggests that the overall connections do not change in the absence of a stimuli.

In the canonical model of ER structures, ER tubules radiate from sheets towards the cell periphery (Westrate et al., 2015), and the two structures are thought not to overlap. However, we observed that tubular structures also reside on the ER sheets themselves (Fig. 2d and Movie 2), which was distinguished by ERnet as seen in Fig. 2d and Movie 3. Like freestanding tubules, they undergo rapid elongation and contractions, which can either lead to new tubular connections (blue arrows), or separations (grey arrows). A subsequent 3D reconstruction of SIM image sections further validated that such tubules are directly attached to the sheets, and are not the result of a projection view artefact (Fig. 2e and Movie 4). Analysis of over 500 cells showed that this phenomenon is a common feature of the ER network (Fig. 2f). Furthermore, we saw that sheet-based tubules form potential contact points for lysosomes. In Extended Data Fig. 4, it is shown that lysosomes play a role to actively guide a tubular structure on sheet domain similar to what has been observed to standard ER-lysosome contact points reported by us recently (Lu et a., 2020).

### ERnet analysis reveals the complex connectivity of the ER tubular network

ERnet can be used to quantify the connectivity of edges and nodes before plotting a corresponding connectivity graph (Fig. 3a). The connectivity graph highlights that the network of the ER largely constitutes of three-way junctions (red nodes, Fig. 3a) while the ER edges are capped with growth ends (green nodes, Fig. 3a).

**Fig. 3:**
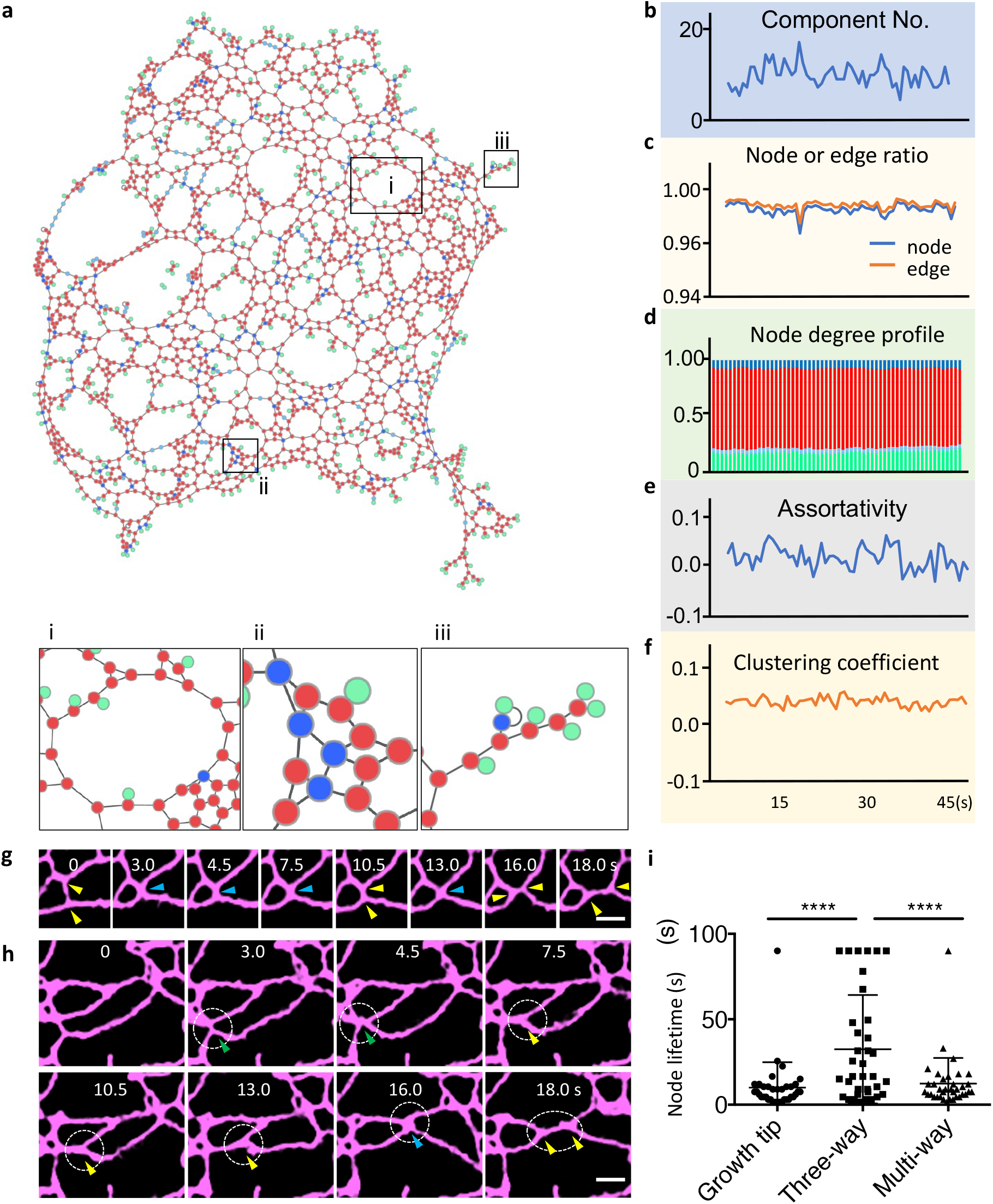
Quantitative analysis by ERnet reveals the complex connectivity of ER tubular network. a. The topology of an ER tubular network is represented by a connectivity graph. i: a polygonal structure organized by three-way junctions and tubules, ii: a representative region of multi-way junctions (dark blue spots), iii: a representative region of ER tubular growth tips (green spots). b-f. Quantitative analysis of the cell shown in (a) over a time window of 45 s. See Source Data Fig. 3b-f. b. Number of components (ER fragments) in time-lapse images. c. Changes of the node or edge ratio over time. d. Quantification of the nodes of different degrees, showing a dominance of third-degree nodes (three-way junctions). Same colour scheme as in (a). e-f. Changes of assortativity and clustering coefficients in time-lapse images. g-h. Examples of transitions between three-way (yellow arrows) and multi-way junctions (yellow arrows: three-way, blue arrows: four-way, green arrows: five-way) junctions. Scale bar: 1 μm. i. Quantification of the lifetime of junctions (nodes) with different degrees. \*\*\*\**P* < 0.0001, Tukey’s one-way ANOVA. n ≥ 20 events per condition from three independent experiments. See Source Data Fig. 3i.

To assess the integrity of the ER, we defined each disconnected ER region as a fragment. As the ER is constantly reshaping, the total number of fragments fluctuates during each recording (Fig. 3b). However, despite these ongoing structural modifications, ERnet reveals that in a typical healthy cell, a single large fragment comprises the majority of all edges and nodes at all times (over 92% of all the 3000 nodes and 95% of all the 2500 edges in the shown example). As quantitative parameters, we defined node and edge ratios (the number of nodes or edges in the largest fragment divided by the total number of nodes or edges, respectively), see Fig. 3c. Per definition, these values range from close to 0 (fully fragmented ER) to 1 (fully connected). Additionally, ERnet quantified the degrees of the ER nodes, *i.e*., how many edges (tubules) connect to each node (junction). As shown in Fig. 3d, three-way junctions are the most abundant and represent 78% of all junction types in this example. Despite the prevailing model of ER morphology, where three-way junctions interconnect to form the whole ER tubular network, ERnet also identified nodes connected with more than three edges (tubules), *i.e.*, multi-way junctions. The presence of multi-way junctions indicates the heterogeneous connectivity of ER tubules that are organised in a higher order of complexity than previously assumed.

Next, the assortativity and clustering coefficients (Fig. 3e and f), that describe connectivity patterns of nodes, were calculated based on the above metrics. The assortativity coefficient measures the tendency of nodes to connect with others of the same degree (Newman 2002) while the clustering coefficient reflects the tendency of nodes to cluster together. Assortativity coefficients range from −1 (fully heterogeneous connectivity, i.e. nodes only connect with those of different degrees) to +1 (fully homogeneous connectivity, i.e. nodes only connect with those of same degree). Clustering coefficients describe another aspect of a node’s connectivity: they measure if the neighbouring nodes of a given node tend to connect to each other, i.e. to cluster. Similarly, for clustering coefficients, 1 describes a perfectly clustered network while 0 refers to no clustering. Fig. 3e shows the ER as a weak assortative network, which suggests a slight tendency of nodes to connect with nodes of the same degree. Additionally, the low clustering coefficients (Fig. 3f) demonstrate a lack of aggregation of nodes and edges in the whole ER of this cell.

To further investigate the structural dynamics of the ER, we tracked the lifetime of multi-way junctions and their transitions from multi-way to three-way junctions. Fig. 3g and h show the rapid transitions between three-way (yellow arrows) and multi-way junctions (blue arrows) driven by ER tubule reshaping. As shown in these cases, the formation of four or five-way junctions need simultaneous connections of more than three tubules at the same junction, which occurs with a lower chance than the formation of a three-way junction that only requires the connection of three tubules. Additionally, any movement of a tubule away from its multi-way junction can lead to the collapse of this junction and the generation of at least two three-way junctions. Therefore, as shown in Fig. 3i, the average lifetime of a multi-way junction is much shorter, *i.e*., less than a third (10.1 s *vs* 30.8 s) of that of a three-way junction.

### Quantitative analysis of ER structures reveals phenotypic characteristics of the ER in stress models

ER morphological defects caused by mutations in genes encoding ER-reshaping proteins or by metabolic perturbations have been linked to a variety of human diseases (Westrate et al., 2015). However, the exact phenotypical ER disruption under these conditions has not yet been sufficiently characterised. Using ERnet, we first analysed the ER morphological defects in stress models mimicking the ER phenotypes in two neurodegenerative diseases, namely Hereditary Spastic Paraplegias (HSPs) and Niemann-Pick disease type C (NPC). The inherited neurological disorder HSPs can be characterised by progressive lower-limb weakness and muscle stiffness, which are caused by mutations in genes encoding ER reshaping proteins such as atlastin (ATL) (Zhao et al., 2001) and protrudin (Mannan et al., 2006). We used ERnet to examine the ER morphology defects in individual cells of different models by measuring two topological features, *i.e*., the degree of ER tubule fragmentation and the heterogeneity in in these tubular connections. Compared with control cells, an ATL knock-out (KO) leads to a collapse of the ER network integrity. Such ER fragmentation was clearly revealed in ATL KO cells by the increasing number of fragments and a 20-fold reduction of the node ratio (99% in control *vs*. 5.4% in ATL KO) (Fig. 4a and Movie 5 and 6). ERnet also highlighted that the lack of ATL significantly altered the connectivity in ER tubular network, as witnessed by a reduced percentage of three-way junctions among all the nodes (26% *vs*. 78% in control) and by the disorganised connectivity (−0.25 in assortativity). These measurements provided quantitative rather than descriptive evidence of ATL’s role in ER tubular network formation, which was previously reported to be crucial for the fusion of ER membranes and, thus, to form continuous networks (Zhao et al., 2001). With these quantitative analyses, we can compare morphological defects caused by different treatments. In another model of HSPs, depletion of protrudin (Extended Data Fig. 5) resulted similarly in ER tubular network fragmentation (305 fragments) (Movie 7) and in disorganised connectivity, however, to a lesser extent. A further metric suitable for the comparison of ER health under different treatments is the size of the ER, which is revealed by the connectivity graph. An ATL KO cell that was more fragmented than a protrudin KD cell suffered from a more severe shrinkage of the ER with a smaller number of nodes and edges (Fig. 4a), indicating that ER membranes may be degraded or recycled in response to stresses. The similar phenotypes observed in both genetic models suggest the connectivity defect in the ER may be a general cause of HSPs.

**Fig. 4:**
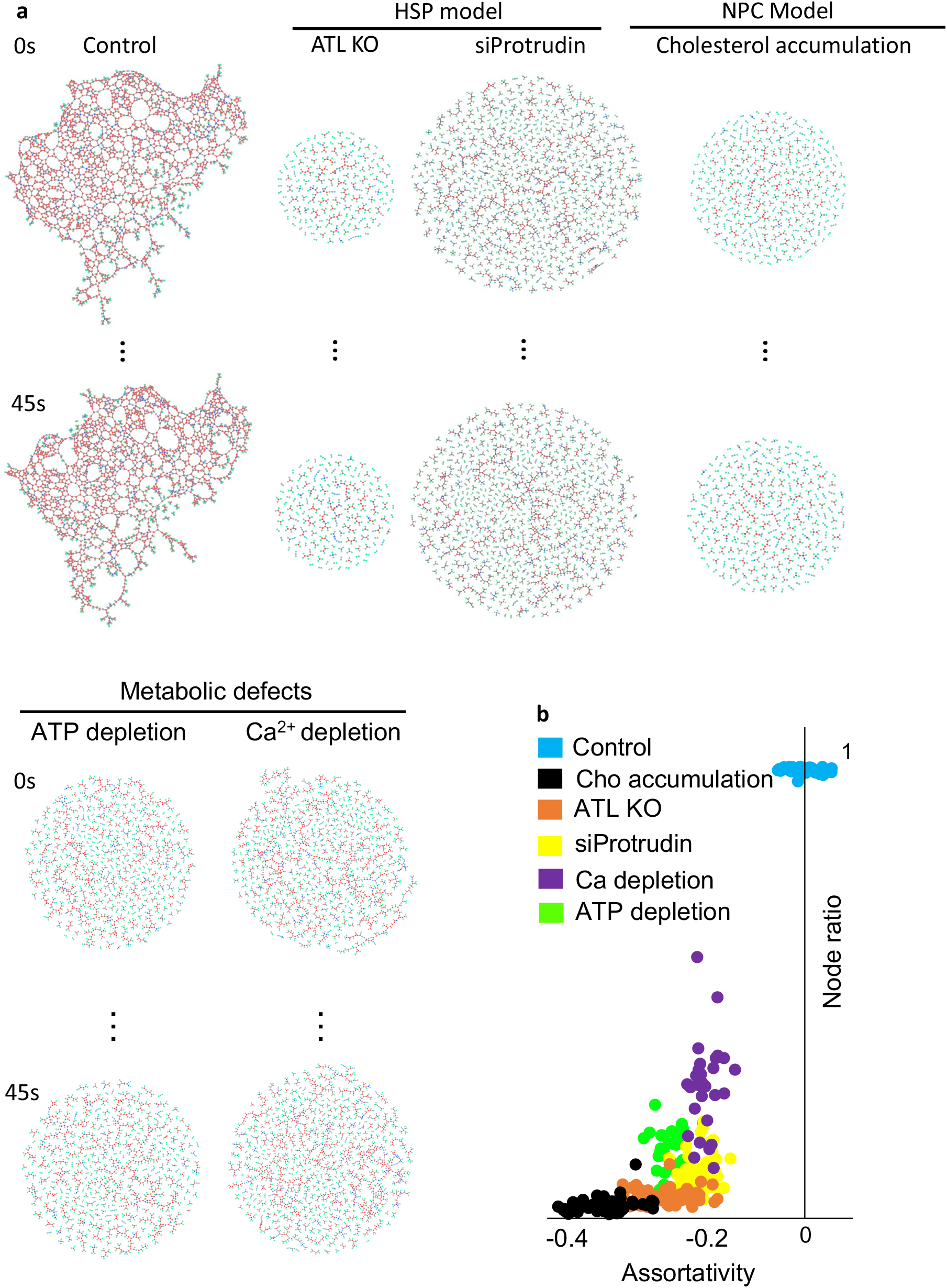
Quantitative analysis of ER phenotypic characteristics in disease associated models. a. Connectivity graphs of ER structures in models mimicking phenotypes of HSPs and NPC and metabolic stress induced by calcium and ATP depletion. Nodes of different degrees are labeled with different colours: green (degree 1), light blue (degree 2), red (degree 3), dark blue (degree >3). b. Topological features of the ER tubular network in above conditions were quantitatively analysed by ERnet. The effects on ER structures from different treatments can be directly visualised and compared by plotting the distribution of node integrating ratio (y axis) and assortativity coefficient (x axis). The analysis of ER phenotype, such as that in ATL KO cells, demonstrated a severe fragmentation and altered connectivity in the numerical data plot. See Source Data Fig. 4b.

Next, we induced cholesterol accumulation in lysosomes by U18666A administration to the cell, which induces a blockage of the cholesterol transfer from lysosomes to the ER in NPC (Ko et al., 2001). The accumulation of cholesterol in lysosomes leads to lysosome deposition in perinuclear regions and, therefore, affects the ER structure and distribution (Lu et al., 2020). However, the exact ER morphological defects have not yet been characterised. ERnet revealed that the ER of U18666A-treated cells features a disassortative network (−0.34) and its low node ratio (3.4%) suggests a highly fragmented structure (Fig, 4a and b, Movie 8), which highlights that lysosomal defects can strongly affect the ER and thus provides us with a useful tool to improve our understanding of organelle dysfunction in NPC.

Finally, we tested performance of ERnet in cells upon ER collapse under metabolic manipulations that significantly affect the overall homeostasis inside the cell. The sequential SIM images showed that the ER largely loses its dynamic reshaping capabilities upon the administration of store-operated calcium entry (SOCE) inhibitor SKF96365 (Merritt et al., 1990) (Movie 9). In the connectivity graph, the ER was largely fragmented and featured as a disassortative network (Fig. 4a and b). Compared with SKF96365, NaN_3_ depletes ATP (McAbee et al., 1987) that supports all the energy consuming processes inside the cell including ER tubule elongation, retraction, and membrane fusion. Therefore, ATP depletion by NaN3 was expected to significantly inhibit the structural dynamics of the ER. ERnet successfully revealed the level of fragmentation of the ER tubular network which resulted from the lack of ATP (Fig. 4a and b, Movie 10); however, such phenotypes were not equivalent to the severe ER defects caused by the depletion of ER reshaping proteins, as the node ratio of ER in ATP depleted cells is nearly 4-fold of that in ATL KO cells (0.19 *vs* 0.05).

Overall, these evaluations highlight the advantages of ERnet to provide quantitative assessments while being sensitive enough to detect the subtle ER morphology changes, especially when it comes to network connectivity, that is required for the investigation of ER-related disease phenotypes.

### Versatility test demonstrates robust performance of ERnet in different cell lines and microscopy techniques

While ERnet has been demonstrated to be suitable for the quantification of ER dynamics in different cell models related to ER stress and diseases, the validation of its robustness and versatility is crucial to ensure its successful application for a wide range of research. Fig. 5 presents the analysis of images obtained using different microscopy techniques including widefield, confocal, and Airyscan microscopy. Even though ERnet’s precision may depend on the spatial resolution of the corresponding images, it performed reasonably well for all imaging techniques with all the tubules and sheets clearly classified and quantified (Source Data Fig. 5). Furthermore, we also performed validation tests for varying cell types commonly used in cell biology research, such as HEK, CHO, SH-SY5Y cells, and primary cultures of hippocampal neurons and glial cells derived from embryonic rats. Although the specific ER phenotypes varied among the cell types, ERnet was able to robustly identify the corresponding tubular and sheet domains and performed subsequent quantitative analyses based on the segmentation (Source Data Fig. 5). The presented reliable segmentations performed on various cell lines and imaging setups further highlight ERnet’s robustness and its precision for the structural analysis of ER networks while providing key metrics suitable to quantify the subtle changes in ER fragmentation and the heterogeneity in tubule connections, crucial for the evaluation of cell healthiness and disease progression.

**Fig. 5:**
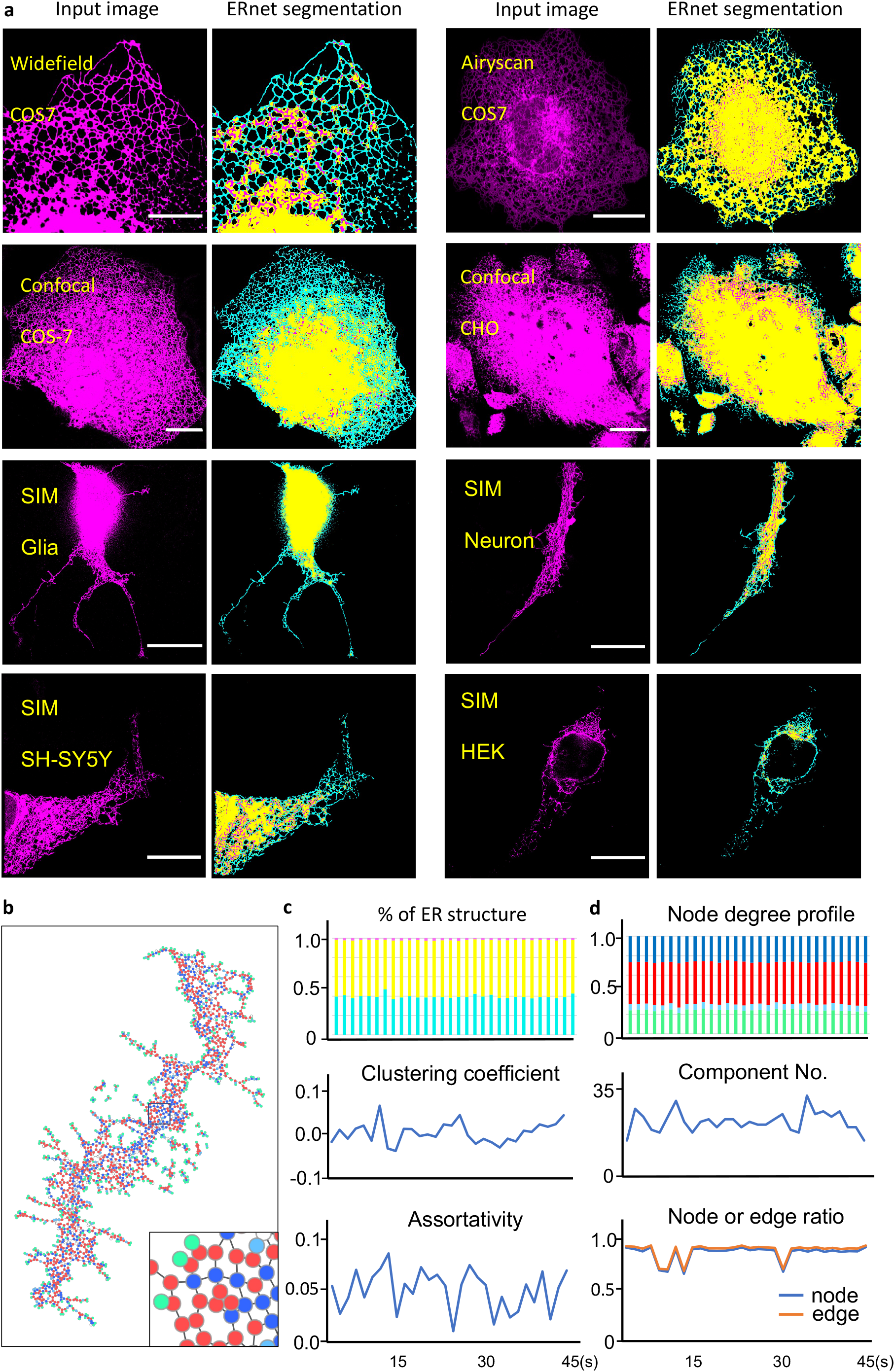
Robust performance of ERnet in versatility test. a. A variety of cell lines with different ER morphologies were imaged by different microscopy techniques to investigate the robustness and versatility of ERnet. ER structures of COS-7, HEK, CHO, SH-SY5Y, primary cultures of hippocampal neurons and glial cells were tested, as well as images acquired by widefield, confocal and Airyscan microscopy. Scale bars: 20 μm. b. The topology of an ER tubular network of the COS-7 cell from the confocal image shown in (a) is represented by a connectivity graph. Nodes of different degrees are labeled with different colours: green (degree 1), light blue (degree 2), red (degree 3), dark blue (degree >3). Bottom right: a zoomed-in region of of the black boxed part in the connectivity, demonstrating the complex connectivity revealed by ERnet from confocal microscopy image. The following analysis of c-f is based on this image data. c. Quantitative analysis of the ER structure of the above image data reveals the topology features of ER tubular network. Top: percentage of the ER tubules (cyan), sheet (yellow), and sheet-based tubules (magenta) of the time-lapse frames. Middle and bottom: changes of assortativity and clustering coefficients in time-lapse images. See Source Data Fig. 5 for c and d. d. Quantitative analysis of the connectivity of the ER tubular network in the above cell. Top: quantification of the nodes of different degrees, showing a dominance of third-degree nodes (three-way junctions). Middle: number of components (ER fragments) in time-lapse images. Bottom: changes of the node/edge ratio over time.

## Discussion

Quantitative cell biology that measures the cellular organelle properties such as shape, position, and mobility provides the basis of analysing the structure and function of organelles in both fundamental and therapeutic research. Here, we introduce ERnet, a versatile tool that performs robust and precise segmentations and permits the quantitative analysis of ER structures in a variety of conditions, including different cell models, cell types and images taken with different microscope techniques. ERnet generates multiple metrics informing on the connectivity of the ER network and permits the quantitative comparisons of ER integrity and structural defects among different stress models. ERnet clearly highlights the fragmented structures and reduced connections of ER networks in stress conditions, which becomes particularly evident in models mimicking phenotypes of HSPs and NPC. While it is difficult and tedious to manually identify and quantify whole ER structures or the fragmented ER pieces of the above models, ERnet provides an automatic and rapid analysis of various phenotypes, which may be used to evaluate disease severity in diagnosis or treatment effects during drug screening.

The high accuracy of ERnet’s semantic segmentation is based on the model design. In contrast to state-of-the-art CNN models commonly used in image segmentation, ERnet is constructed in a Vision Transformer architecture that outperforms CNNs with higher accuracy in image classification tasks but with four times fewer computational resources (Dosovitskiy et al., 2020; Paul and Chen 2021). Another advantage of our design is its capability for temporal domain analyses of objects from sequential images. We also integrated two attention mechanisms: multi-head self-attention and channel attention into the Transformer architecture. These mechanisms greatly enhance the learning ability of ERnet in classifying ER structures in the spatio-temporal domain. While machine learning methods have previously been implemented to reconstruct ER structures based on electro-microscopy images (Liu et al., 2019) and to identify ER stress marker-whorls (Guo et al., 2022), ERnet can be applied for video-rate image segmentation and the analysis of live cells, thus, further extending the deep learning toolbox for biomedical research.

By applying ERnet, we characterised the structural features of the dynamic ER network. First, we found that the dominance of three-way junctions is a necessity to produce a continuous ER network which can spread throughout the cell and, in addition to the prevalence of three-way junctions, it has been observed that a healthy ER contains approximately 20% of multi-way junctions (degree > 3). In contrast, all the stress manipulations of ER morphology, including models of HSPs and NPC, resulted in the fragmentation of ER structures to varying extents (Fig. 4). Although the ER fragmentation may be easily visualised in images, it is difficult to evaluate the severity of fragmentation caused by different treatments and even harder to compare based on descriptive imaging data. ERnet not only demonstrates the degree of fragmentation, but also analysed this morphological defect from different angles with a list of metrics. Therefore, we can have a quantitative and comprehensive understanding of the ER phenotype and a reliable comparison of treatments by plotting the numerical data informing us on the level of ER fragmentation and connectivity in a same framework (Fig. 4b). We showed an example of multi-parameter analysis of ER in single cells in sequential frames, demonstrating the consistency of the phenotype during the recording (Fig. 4b). This consistency is more prominent in the population level, as the data point to different cells under the same condition grouped together and separated from the data from other conditions (Extended Data Fig. 6). This demonstrates that ERnet is suitable to detect and measure phenotypic characteristics of the ER in different cell populations. All these provide a powerful tool to investigate potential therapies for ER associated diseases.

Another key advantage of deep learning-based image processing is their ability to drive novel biological observations. Since ERnet is sensitive to structural features, our model was able to identify sheet-based tubules. These ER components share similar structures and dynamics with the tubules that radiate from the sheet domains towards the periphery of the cell, however, their position in the sheet domain greatly extends the coverage of the tubular ER towards the cell centre and even close to the nucleus. Finally, the observed sheet-based tubules’ close contact with lysosomes might permit beneficial material exchange and structure regulation as lysosomes are one of the cell’s sensing hubs. How the sheet-based tubules are regulated in both physiological and pathological conditions will be an important question for future studies.

We believe our work demonstrates an efficient tool for precise structure segmentation and multi-parameter analysis of ER phenotypes. Its user-friendly graphical interface and automatic batch processing can save a significant amount of manual curation in imaging annotation and, therefore, speed up ER associated disease research and therapeutic screenings. In the future, the integration of ERnet with other organelle analysis tools, such as methods for lysosomes and mitochondria characterisations, will open the door to quantitative and comprehensive investigations of multi-organelle interactions and their roles in the development, degeneration, and ageing of cells.

## Acknowledgments

We thank Dr Ana Isabel Fernández Villegas and Yuqing Feng for helping with the cell culture. We thank Dr Edward Ward for helping with the image processing. We thank Prof. Junjie Hu (Chinese Academy of Sciences, China) for giving us the ATL KO cell line.

## Author contributions

ML designed, conducted, and interpreted experiments, and wrote the article. ML and C.N.C. developed the whole pipeline of ERnet. C.N.C. developed the core model of ERnet. J.W. provided the graph-theory based analysis of ERnet. T.K. supported the versatility test. NL, KS, E.A., P.L., A.L. and G.S.K.S. gave advice and edited the article. C.F.K. supervised the research, coordinated the study, and wrote the article.

## Data availability

All data needed to evaluate the conclusions in the paper are present in the Source Data files. Additional data related to this paper may be requested from the corresponding authors.

## Code Availability

The ERnet model is written in Python. The software and Colab versions of ERnet are also freely available online through GitHub at https://github.com/charlesnchr/ERnet-v2.

## Methods

### Cell culture

COS-7 cells were purchased from the American Type Culture Collection (ATCC). COS-7 cells were grown in T75 or T25 flasks or six-well plates by incubation at 37°C in a 5%CO_2_ atmosphere. Complete medium for normal cell growth consisted of 90% Dulbecco’s modified Eagle’s medium (DMEM), 10% fetal bovine serum (FBS) and 1% streptomycin. Cells were kept in logarithmic phase growth and passaged on reaching 70 to 80% confluence (approximately every 3 to 4 days). Medium was changed every 2 or 3 days. For structured illumination microscopy (SIM) imaging experiments, COS-7 cells were plated onto Nunc Lab-Tek II Chambered Coverglass (Thermo Fisher Scientific, 12-565-335) to achieve ~70% confluence before transfection.

COS-7 cells were transfected with mEmerald-Sec61b-C1 (Addgene #90992, gifted by Jennifer Lippincott-Schwartz, Janelia Research Campus) as indicated with Lipofectamine 2000 according to the manufacturer’s protocol 24 to 48 hours before imaging. Cells were stained with SiR-Lysosome at 1 μM for 4 hours before imaging. Cells were imaged in a microscope stage top micro-incubator (OKO Lab) with continuous air supply (37°C and 5% CO_2_). Cells were treated with U18666A (662015, Sigma) at 10 μM for 24 hr to block cholesterol transfer from lysosomes to ER before imaging. Cells were treated with SKF-96365 (S7809, Sigma) at 100 μM for 3 hr to deplete Calcium before imaging. Cells were treated with NaN_3_ (0.05% w/v) and 2-deoxy-glucose (20 mM) for 2 hr to deplete ATP before imaging. SH-SY5Y cells were cultured and images as previously described (Michel et al., 2014). HEK cells were cultured and imaged as previous described (Lu et al., 2019). ATL KO model was gifted by Prof. Junjie Hu, Chinese Academy of Sciences, China. CHO-K1 cells were purchased from ATCC and were cultured in Ham’s F-12 Nutrient Mixture medium supplemented with 10% FBS, 2 mM L-Glutamine and 100 U/mL Penicillin-Streptomycin (Pen/Strep). Cells were transfected with pFLAG_ER mCherry (Avezov et al., 2015). U2OS cells were purchased from ATCC and were cultured in DMEM supplemented with 10% FBS, 2 mM L-Glutamine and 100 U/mL Pen/Strep. Cells were transfected with pFLAG_ER mCherry (Avezov et al., 2015).

### siRNA transfection and Western

#### blot

Protrudin were depleted using SMARTpool: ON-TARGETplus, Dharmacon. Negative siRNA control (MISSION siRNA Universal negative control) was purchased from Sigma-Aldrich. COS-7 cells were plated in both glass-bottom Petri dishes (for imaging) and six-well plates (for Western blot validation). Cells were transfected with 20 nM siRNA oligonucleotides and 20 nM negative control siRNA using Lipofectamine RNAiMax (Thermo Fisher Scientific) according to the manufacturer’s protocol. After 6 hours of siRNA transfection, the cells were washed and the medium was replaced with complete culture medium. Twenty-four hours after the siRNA transfection, cells were transfected with plasmid DNA indicated in Results using Lipofectamine 2000 (Invitrogen). On the day of imaging, cells were stained with Sir-Lysosome. Cells in glass Petri dishes were imaged 24 hours after DNA transfection.

Cells in six-well plates were harvested for Western blot validation 72 hours after siRNA transfection. Protein concentration was measured using a bicinchoninic acid (BCA) protein assay kit. Immunoblotting was performed by standard SDS–polyacrylamide gel electrophoresis/Western protocols. Primary antibody concentrations were as follows: anti-Protrudin at 1:5000; GAPDH (glyceraldehyde-3-phosphate dehydrogenase) at 1:30,000; tubulin at 1:5000. Secondary antibodies (Sigma-Aldrich) were used at 1:3000 for all rabbit antibodies and for all mouse antibodies. The signal was detected with SuperSignal West Pico Chemiluminescent Substrate.

### Widefield and Structured illumination microscopy

SIM imaging was performed using a custom three-color system built around an Olympus IX71 microscope stage, which we have previously described (Young et al., 2016). Laser wavelengths of 488 nm (iBEAM-SMART-488, Toptica), 561 nm (OBIS 561, Coherent), and 640 nm (MLD 640, Cobolt) were used to excite fluorescence in the samples. The laser beam was expanded to fill the display of a ferroelectric binary Spatial Light Modulator (SLM) (SXGA-3DM, Forth Dimension Displays) to pattern the light with a grating structure. The polarization of the light was controlled with a Pockels cell (M350-80-01, Conoptics). A 60×/1.2 numerical aperture (NA) water immersion lens (UPLSAPO 60XW, Olympus) focused the structured illumination pattern onto the sample. This lens also captured the samples’ fluorescent emission light before imaging onto an sCMOS camera (C11440, Hamamatsu). The maximum laser intensity at the sample was 20 W/cm^2^. Raw images were acquired with the HCImage software (Hamamatsu) to record image data to disk and a custom LabView program (freely available upon request) to synchronize the acquisition hardware. Multicolour images were registered by characterising channel displacement using a matrix generated with TetraSpeck beads (Life Technologies) imaged in the same experiment as the cells. COS-7 cells expressing mEmerald-Sec61b-C1 (ER marker) and stained with SiR-Lysosome (lysosome marker) were imaged by SIM every 1.5 s (including imaging exposure time of both channels) for 60 frames.

### Reconstruction of the SIM images with LAG SIM

Resolution-enhanced images were reconstructed from the raw SIM data with LAG SIM, a custom plugin for Fiji/ImageJ available in the Fiji Updater. LAG SIM provides an interface to the Java functions provided by fairSIM (Müller et al., 2016). LAG SIM allows users of our custom microscope to quickly iterate through various algorithm input parameters to reproduce SIM images with minimal artifacts; integration with Squirrel (Culley et al., 2018) provides numerical assessment of such reconstruction artifacts. Furthermore, once appropriate reconstruction parameters have been calculated, LAG SIM provides batch reconstruction of data so that a folder of multicolour, multi-frame SIM data can be reconstructed overnight with no user input.

### AiryScan imaging

AiryScan imaging was performed using a LSM 880 confocal microscope (Zeiss). A Zeiss Plan-Apochromat 63×/1.40 DIC M27 Oil objective was used. For visualisation of ER structure, ER mCherry was excited by a diode-pumped solid-state (DPSS) 561 nm laser (1% intensity) and detected using the AiryScan detector. Bit depth was set at 16 bits. Using the Fast-Airyscan mode, live-cell time-lapse images were acquired every 1 second (60 frames) with an image size of 1364 × 1244 pixels. Cells were kept in a controlled environment (37°C, 5% CO_2_) during imaging. Following acquisitions, images were deconvoluted using the Airyscan processing. Image processing was performed in software ZEN 2.3 SP1 FP3 (black) (ver.14.0.25.201).

### Confocal Imaging

A part of confocal imaging was performed using a STELLARIS 8 confocal microscope (Leica). A HC PL APO CS2 63x/1.40 OIL objective was used. For visualisation of ER structure, ER mCherry was excited by 587 nm of white light laser (WLL) with 3% intensity and detected using the HyD S3 detector (detection range: 592-750 nm). Bit depth was set at 16 bits. Live-cell time-lapse images were acquired every 1.5 seconds (90 frames) with an image size of 512 × 512 pixels. Cells were kept in a controlled environment (37°C, 5% CO_2_) during imaging.

### ERnet construction

For the segmentation of the sequential endoplasmic reticulum (ER) images, a spatio-temporal shifted window vision transformer neural network is trained and used. The proposed model is inspired by the previous models Vision Transformer (Dosovitskiy et al. 2020), its more efficient shifted window variant Swin (Liu et al. 2021), with its extension for video classification Video Swin (Liu et al. 2021a), and adaption to image restoration SwinIR (Liang et al. 2021). Swin introduced the inductive bias to self-attention called shifted window multi-head attention (SW-MSA) which can be compared to the inductive bias inherent in convolutional networks. SwinIR introduced residual blocks to the Swin transformer to help preserve high-frequency information for deep feature extraction. The Video Swin transformer extended the SW-MSA to three dimensions, such that spatio-temporal data can be included in the local attention for the self-attention calculation. Further to this, the success of the channel attention mechanism (Zhang et al. 2018) inspired the inclusion of this other inductive bias in addition to 3D local self-attention following the SW-MSA approach.

The inputs to the model have the dimension *T* × *H* × *W* × *C*, where *T* is 5 for ERnet (5 adjacent temporal frames) and *C* is 1 (grayscale inputs). A shallow feature extraction module in the beginning of the network architecture, shown in Fig. 1, projects the input into a feature map, *F*_0_, of *T* × *H* × *W* ×*D* dimension, where the embedding dimension, *D*, is a hyperparameter. The feature map is passed through a sequence of residual blocks denoted Window Channel Attention Block (WCAB)

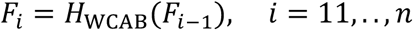

Inside each WCAB is a sequence of Swin Transformer Layers (STLs), in which multi-head self-attention is calculated using local attention with shifted window mechanism. Inputs to STL layer is partitioned into 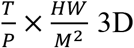 tokens of *P* × *M*^2^ × *D* dimension. For a local window feature, 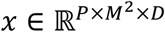, query, key and value matrices, 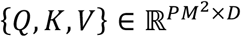, are computed by multiplication with projection matrices following the original formulation of transformers. Attention is then computed as

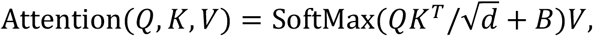

where 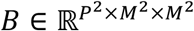 is a relative positional bias found to lead to significant improvements in classification performance. STLs are joined in a way similar to the residual blocks, although the use of SW-MSA is alternated with a version without shifted windows, W-MSA, ensuring that attention is computed across window boundaries, which would not have been the case without SW-MSA.

After the final STL, the *m*-th layer, in a WCAB, a transposed 3-dimensional convolutional layer is used to project the 3D tokens back into a *T* × *H* × *W* × *D* feature map, *F_i,m_*. A channel attention module is then used on *F_i,m_* to determine the dependencies between channels following the calculation of the channel attention statistic. The mechanism works by using global adaptive average pooling to reduce the feature map to a vector which, after passing through a 2D convolutional layer, becomes weights that are multiplied back onto *F_i,m_* such that channels are adaptively weighed. A residual is then obtained by adding a skip connection from the beginning of the *i*-th WCAB to prevent the loss of information, *i.e*., low-frequency information, and the vanishing gradient problem. A fusion layer combines the temporal dimension and the channel dimensions. For the final upsampling module, we use the sub-pixel convolutional filter to expand the image dimensions by aggregating the fused feature maps.

The model is trained by minimising a multi-class cross-entropy loss function

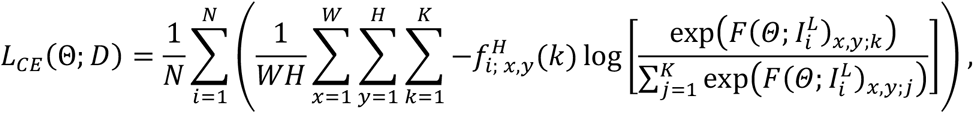

where *k* and *j* are iterators over a total of *K* unique classes, and 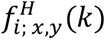 is a function equal to 1 if the target class for the pixel at (*x, y*) of the *i*^th^ image is *k* and equal to 0 otherwise. In this paper, we study the segmentation of background, tubules, sheets, and sheet-based tubules and, therefore, *K* = 4 in the equation above.

The training data is obtained by acquiring experimental data using structured illumination microscopy (SIM). A total of 20 sequential stacks of different samples are acquired, where each stack consists of 60 SIM images reconstructed with ML-SIM. The super-resolved SIM outputs are then segmented by manually finetuning a random forest model in the Weka plugin for ImageJ on an image-by-image basis.

### Network analysis methods

To quantify the structural changes in the ER, methods from network analysis are applied (Boccalettti et al., 2006; Costa et al., 1987). We represent the ER structure of tubules through an undirected and unweighted graph. All tubule junctions are represented by nodes, and the tubules by edges.

Networks are built in a python routine and their metrics are measured through the python package *graph-tool* (Peixoto 2014) and *network x* (Hagberg et al., 2008). We measure the size of the network through the number of nodes: *N*, and edges: *E*, within the system. The number of edges attached to one node is called the nodes degree: *k*, and the distribution of the degrees is one of the most fundamental parts of the analysis of network structures.

To quantify the structural arrangements of the ER, we focus on primary network connectivity metrics. Firstly, we measure the network density, *d*, between nodes and edges (see Eq. (2)). Other metrics that describe the network connectivity are the global clustering coefficient (see Eq. (2)) and the network assortativity (see Eq.(3)). The global clustering coefficient describes the tendency of the network to build triangles, by relating triplets to each other. Three nodes connected to each other through three edges are a *closed triplet*, while three nodes connected to each other through two edges are called an *open triplet* (Newman 2003). The network assortativity describes the likelihood of nodes connecting with nodes of similar properties; here specifically, as is common, a node degree. Assortative mixing is contrasted to disassortative mixing where nodes tend to connect to others of dissimilar properties (Cimini et al., 2019). The assortativity coefficient, *r*, is described in Eq.(3), where *e_i,j_* is the fraction of edges linking a node with type *i* to nodes of type *j*, *a_i_* is the sum over *e_i,j_* for all *j* and *b_i_* is the sum over *e_i,j_* for all *i*. An assortativity coefficient of *r* = 0 indicates no mixing preference, whereas positive values indicate assortative and negative values disassortative tendencies.

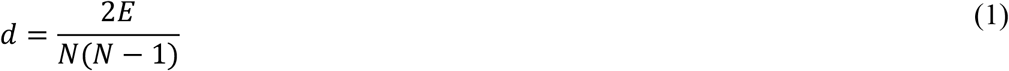

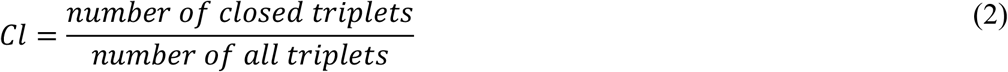

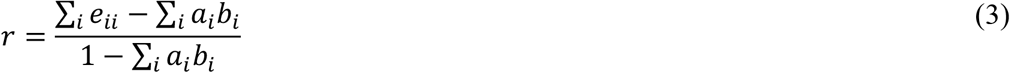

Additionally, we include macroscopic network arrangements by counting the number of network components. Networks may be entirely connected or composed of many distinct components (Albert 2005). For networks evolving over time, network components outline merging or splitting behaviour. In networks with many components, the most characteristic topological features are often exhibited in the largest component (Storgatz 2001).

### Data visualization

Videos of time-lapse imaging and analysis were performed using Fiji (NIH). The connectivity graphs in the figures are re-plotted by a Python module named “connectivity graph.py”. Instructions of using this module is provided inside the file.

### Statistical analysis

Statistical significance between two values was determined using a two-tailed, unpaired Student’s *t* test (GraphPad Prism). Statistical analysis of three or more values was performed by one-way analysis of variance with Tukey’s post hoc test (GraphPad Prism). All data are presented as the mean ± SEM; \**P* < 0.05, \*\**P* < 0.01, \*\*\**P* < 0.001, and \*\*\*\**P* < 0.0001. Statistical parameters including the exact value of *n*, the mean, median, dispersion and precision measures (mean ± SEM), and statistical significance are reported in the figures and figure legends. Data are judged to be statistically significant when *P*< 0.05 by two-tailed Student’s *t* test. In the figures, asterisks denote statistical significance as calculated by Student’s *t* test (\**P* < 0.05, \*\**P* < 0.01, \*\*\**P* < 0.001, and \*\*\*\**P* < 0.0001).

---

**Extended Data Fig. 1:**
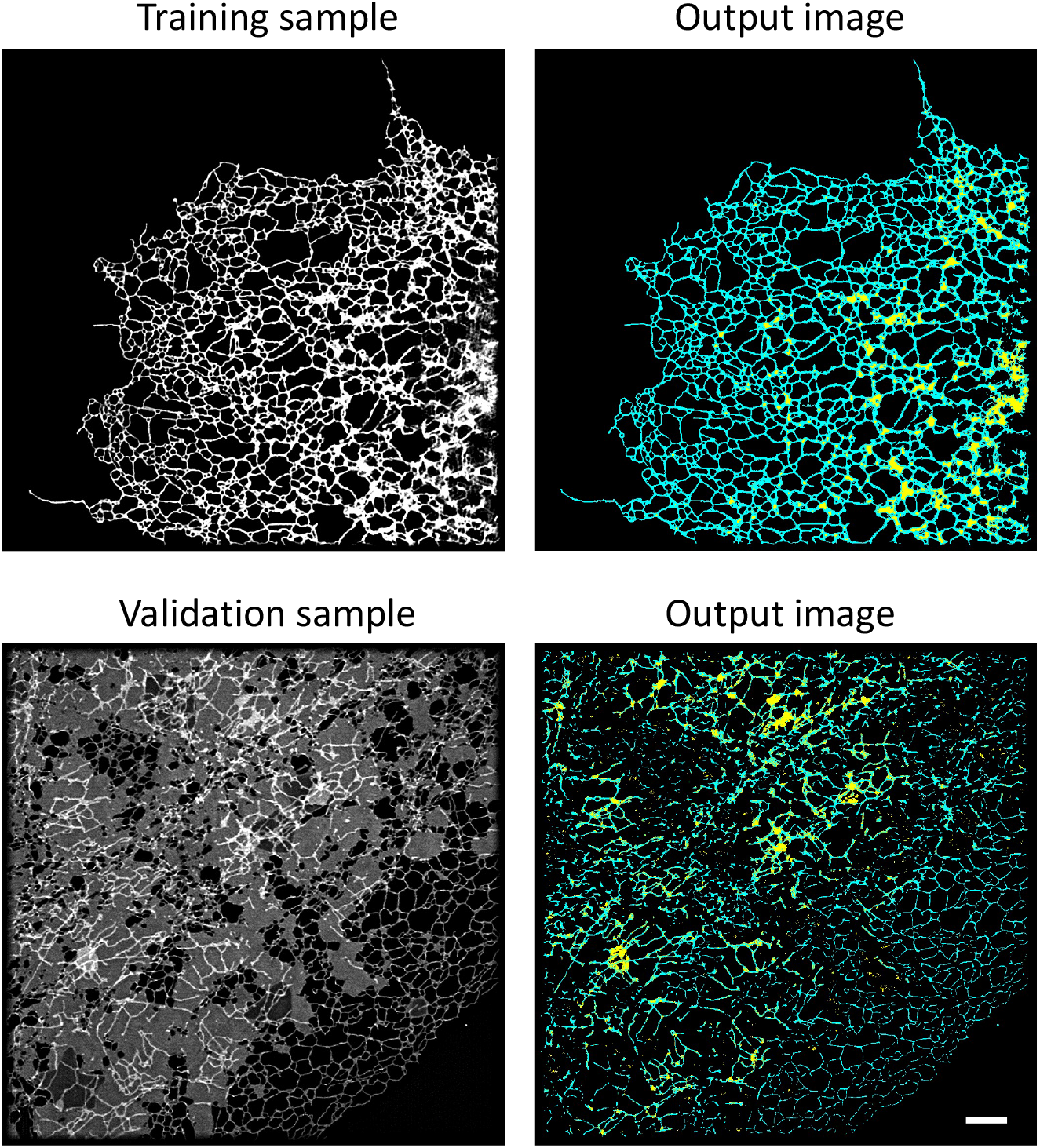
A test Weka trainable segmentation with different input data. Top left: An input image was used to train a classifier of Weka Trainable Segmentation. Top right: The tubules (red) and sheet (green) can be clearly classified after segmentation. Bottom left: a new image was applied to the trained classifier shown above. Bottom right: segmentation result of the new input data. Scale bars 5 μm.

**Extended Data Fig. 2:**
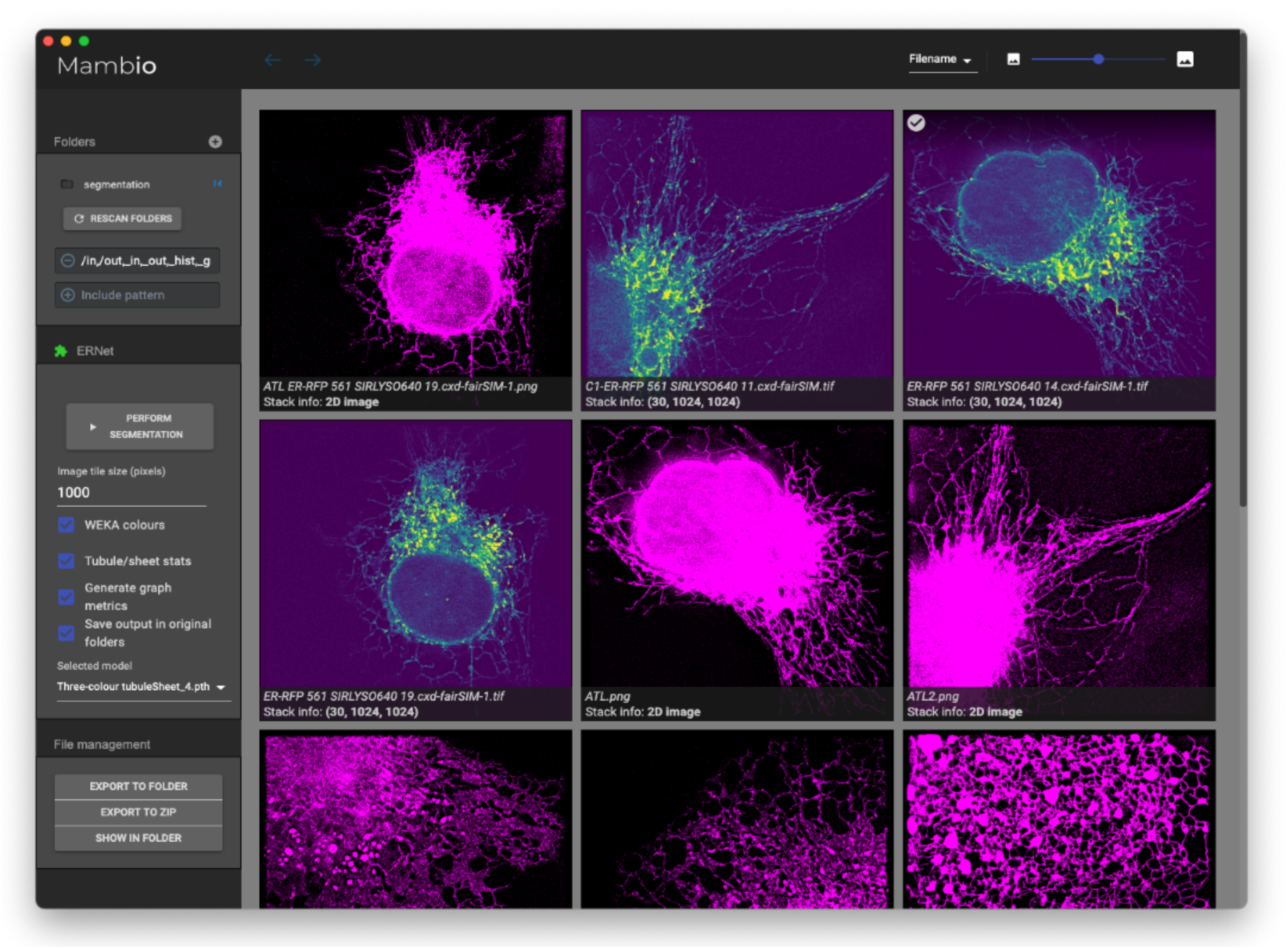
ERnet graphical user interface. Left part of the interface shows the path of input and output images. Bottom left: options of the analysis provided by ERnet. Right part of the interface shows the input images (magenta) and segmented results.

**Extended Data Fig. 3:**
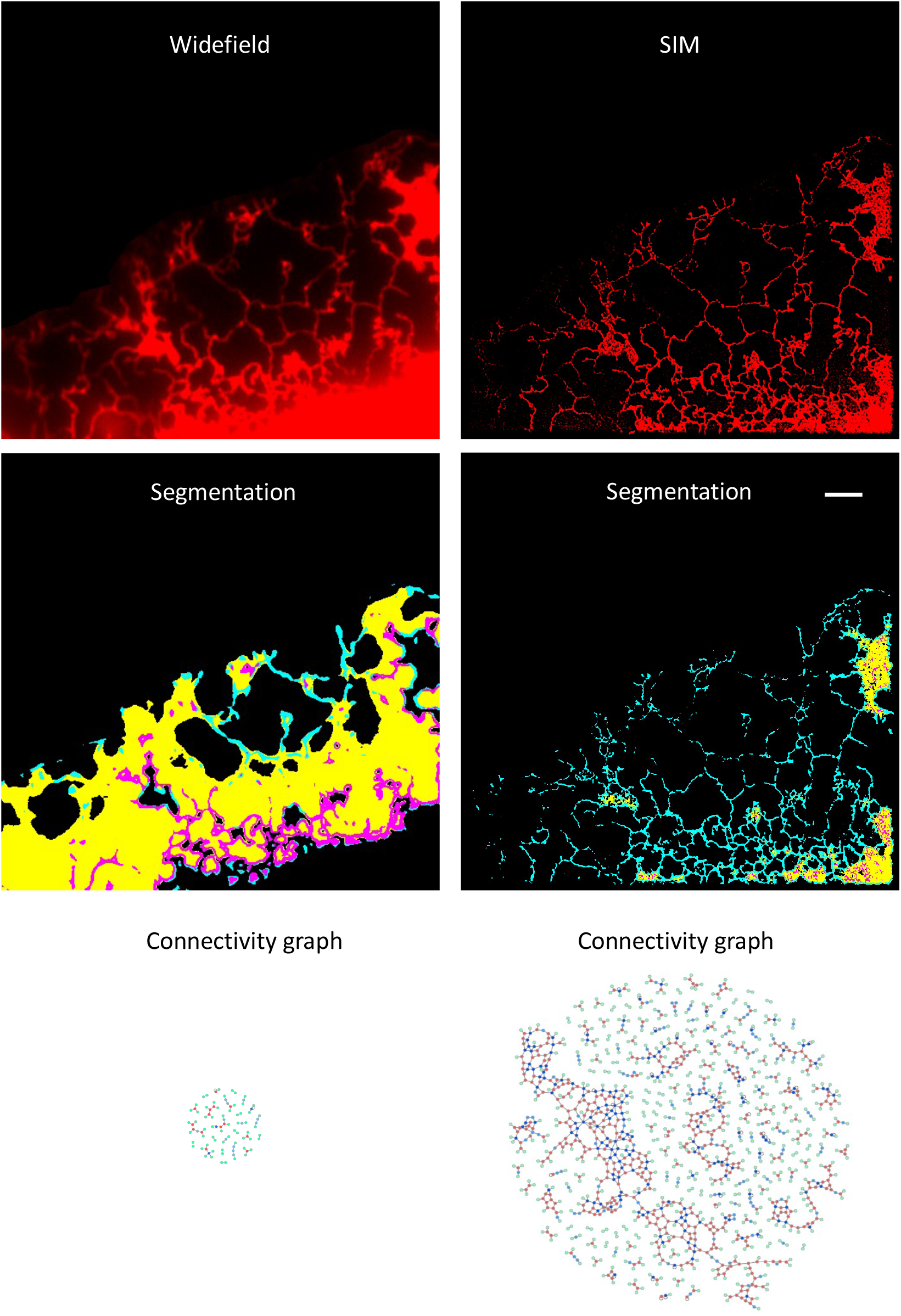
High spatial resolution and signal-to-noise-ratio in SIM image compared with widefield image. Top panel: widefield and SIM images of an ER (red) in a COS-7 cell expressing mEmerald-sec61b-C1. Middle panel: segmentation performed by ERnet of the above images. Bottom panel: connectivity graph plotted based on the topology data quantified from the above segmentation. Scale bar: 5 μm

**Extended Data Fig. 4:**
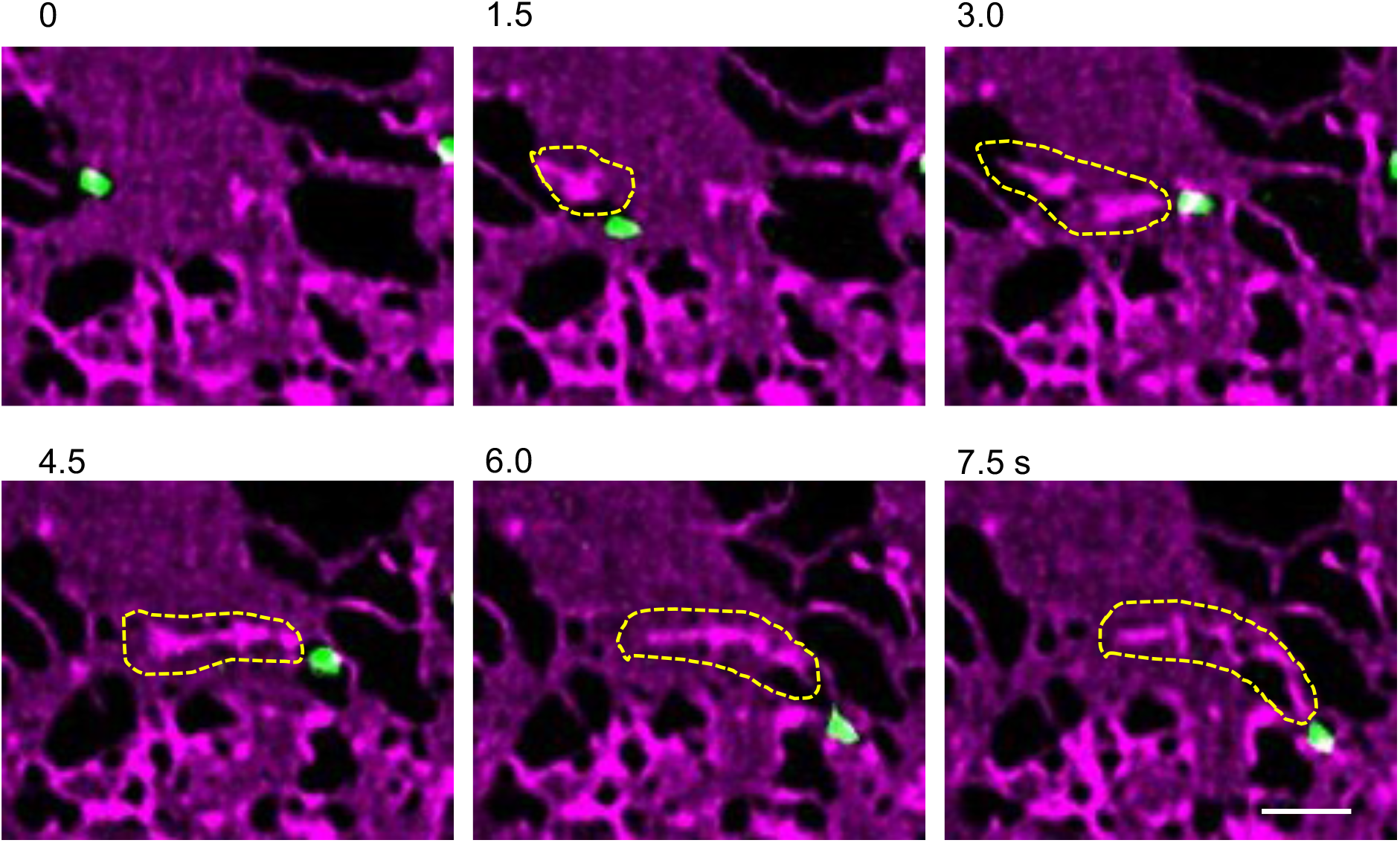
Coupled motion of lysosome and sheet-based tubule indicating inter-organelle contacts between them. Time-lapse SIM images show a lysosome leading a tubule while sliding on a sheet, indicating the motion of this sheet-based tubule is induced by the motile lysosome. The sliding sheet-based tubules are highlighted by yellow dashed line. Scale bar: 2 μm.

**Extended Data Fig. 5:**
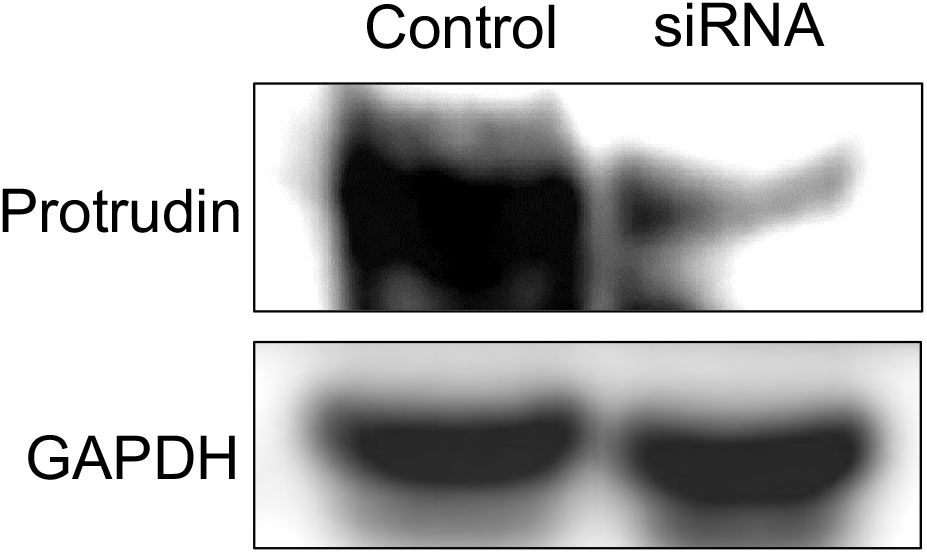
Western blot validation of Protrudin depletion.

**Extended Data Fig. 6:**
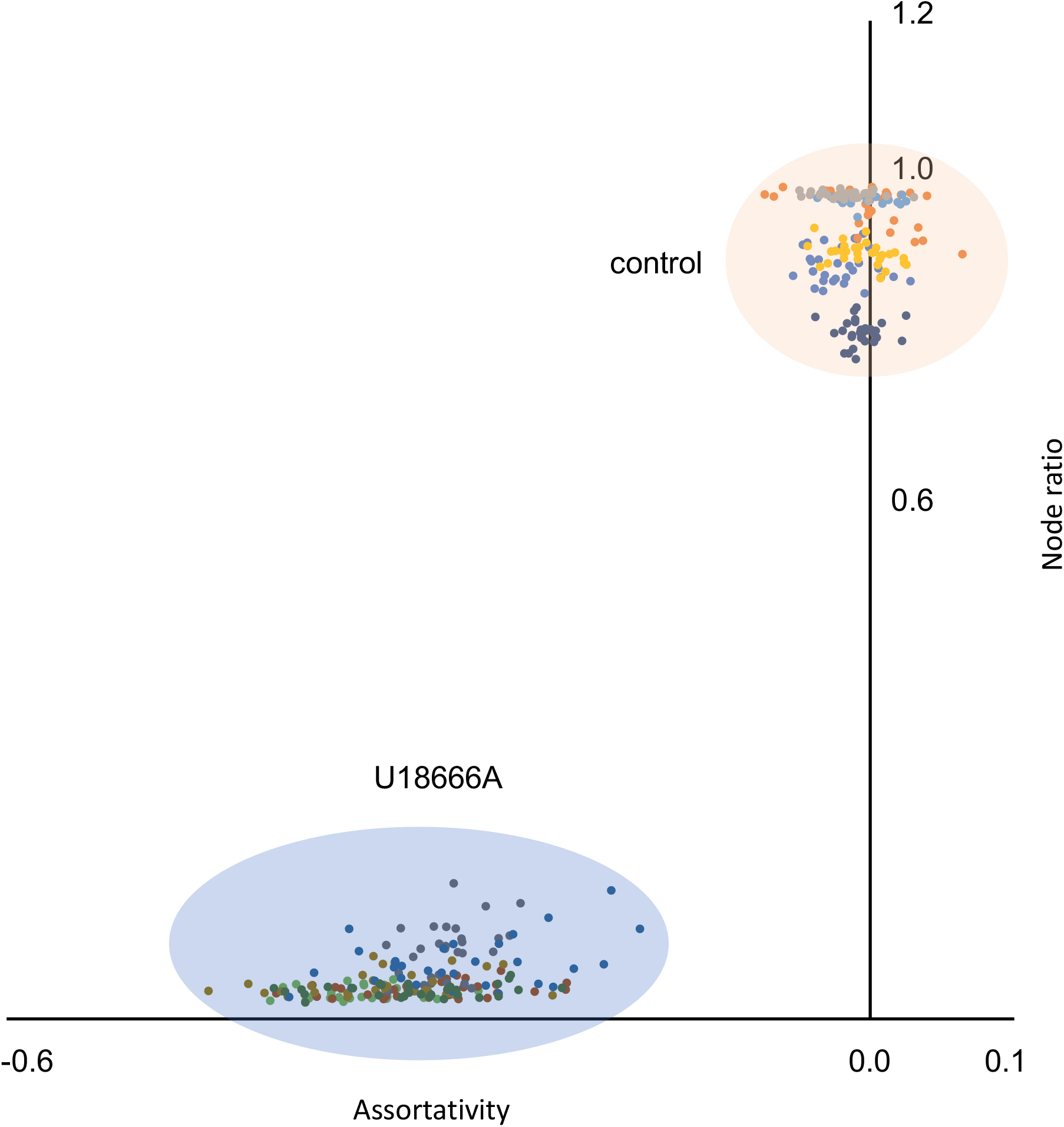
Analysis by ERnet revealing the phenotype consistency in the cell population. Data from the same cell are plotted in the same colour. Time-lapsed SIM images (30 frames, 1.5s/frame for all the data points) of ER structure in each single cell were segmented and analysed by ERnet. The light orange and blue backgrounds suggest the grouped distribution of the data points from the same condition. See Source Data Extended Data Fig. 6.

